# Experience-dependent inhibitory plasticity is mediated by CCK+ basket cells in the developing dentate gyrus

**DOI:** 10.1101/2020.04.30.071126

**Authors:** Ting Feng, Christian Alicea, Vincent Pham, Amanda Kirk, Simon Pieraut

## Abstract

Early postnatal experience shapes both inhibitory and excitatory networks in the hippocampus. However, the underlying circuit plasticity is unclear. Using an enriched environment (EE) paradigm, we assessed the circuit plasticity of inhibitory cell-types in the hippocampus. We found that cholecystokinin (CCK)-expressing basket cells strongly increased somatic inhibition on the excitatory granular cells (GC) following EE while another pivotal inhibitory cell-type, parvalbumin (PV)-expressing cells did not show changes. By inhibiting activity of the entorhinal cortex (EC) using a chemogenetic approach, we demonstrate that the projections from the EC is responsible for the developmental plasticity of CCK+ basket cells. Our measurement of the input decorrelation by DG circuit suggests that EE has little effect on pattern separation despite of the altered CCK+ basket cell circuit. Altogether, our study places the activity-dependent remodeling of CCK+ basket cell innervation as a central process to adjust inhibition in the DG, while maintaining the computation in the circuit.

## Intro

Early postnatal development is a critical period during which most synapses are formed. At this time, early sensory stimulations are critical for proper circuit maturation and can have long lasting deleterious effect on sensory processing as well as on complex cognitive functions when compromised (Cancedda, 2004; Kalogeraki, Pielecka-Fortuna, & Löwel, 2017; Nabel & Morishita, 2013; Sale et al., 2004). Neuronal activity plays a critical role in regulating experience-dependent plasticity and in fine-tuning neural circuits originally assembled under the control of genetic programs. Pre-weaning enrichment (PE), i.e. when pups are raised with their mother from birth to weaning time, offers a valuable model to study the cellular mechanisms underlying experience-dependent remodeling of nascent neural circuits. Studies using PE found significant alterations in the synaptic network of the dentate gyrus (DG) in the hippocampus. Electrophysiology data demonstrate an increase in excitatory and inhibitory drives in juvenile mice raised in EE during this pre-weaning period (Hosseiny et al., 2015; Liu, He, & Yu, 2012). At the structural level, increased spine density is consistently found on the dendrites of excitatory GCs suggesting that the increase in the excitatory drive is due to an increased number of synapse formed by the afferents from the entorhinal cortex layer 2 (EC2) onto GC dendrites (Hosseiny et al., 2015; Liu et al., 2012). However, the underlying cellular and circuit mechanism of the increased inhibitory drive is not known. Particularly, the involvement of specific population of inhibitory synapses in this plasticity has not been explored. This question is particularly challenging because diverse classes of inhibitory interneurons innervate the DG. The well-established inhibitory cell-types that have been described in the cortical and other hippocampal circuits have also been characterized in the DG (Hosp et al., 2014; Pelkey et al., 2017). Interestingly, each inhibitory cell-type projects their axons to distinct laminae where they target specific subdomains along the somatodendritic axis of principal excitatory GCs. This unique connectivity rule enables pattern separation, a process by which the rich and overlapping multimodal inputs from the entorhinal cortex are decorrelated into a sparse and specific output pattern (Barak et al., 2019; Bel, Pardi, Ogando, Schinder, & Marin-Burgin, 2015; Espinoza, Guzman, Zhang, & Jonas, 2018; C.-T. T. Lee et al., 2016; Madar, Ewell, & Jones, 2019b; Szabo et al., 2017). This computation is critical for information coding in the hippocampus and for memory indexation and retrieval (Deng, Mayford, & Gage, 2013; Kesner, Kirk, Yu, Polansky, & Musso, 2016). Understanding how GABAergic synaptic network is shaped by postnatal experience can inform on how the computations in the hippocampus are maintained during this period of dramatic connectivity change. Do all inhibitory cell-types indistinctly form more synapses when animals are raised in EE, or are there specific cell-types responsible for the increased inhibitory drive?

To address these questions, we measured the synaptic densities from the distinct population of inhibitory neurons in PE mice. We found that enriched postnatal experience strongly impacts the number of cholecystokinin (CCK)-expressing basket cell inputs on GC soma. In contrast, innervation from parvalbumin (PV)-expressing cells was unaffected. By combining our enrichment paradigm with a chemogenetic approach, we demonstrate that projection neuron activity in EC is instructive for this structural plasticity. Furthermore, we measured the decorrelation function in GC using slice physiology and found that pattern separation appeared largely preserved despite the strong remodeling of the synaptic network. Altogether, our work brings new insight into how specific inhibitory cell types undergo structural rearrangement in response to the changes in cortical activity. Furthermore, we show that this experience-dependent remodeling of dentate connectivity has a limited impact on pattern separation performed by individual GC.

## Results

### Pre-weaning enrichment increases inhibitory and excitatory drive in the DG

To investigate how early postnatal experience impacts the synaptic network in the hippocampus, mice were raised in a standard laboratory cage or in a large cage containing toys, a hamster wheel and diverse objects that were changed every other day (**Figure 1A**). In these enriched environment (EE) cages, pups were raised for 21 days with their mothers and an additional female to increase nurturing. Miniature inhibitory and excitatory postsynaptic currents (mIPSC and mEPSC) were then recorded from the excitatory granular cells (GCs) in the DG using acute brain slices (**Figure 1B**). We found a significant increase in the frequency of miniature excitatory postsynaptic currents (mEPSCs) (**Figure 1C)** (SH: 1.82 ± 0.21Hz, PE: 2.40 ± 0.25 Hz, p = 0.025), with no significant changes in mEPSC amplitudes (SH: 7.299 ±0.446 pA, PE: 6.74 ± 0.51 pA, p = 0.44). Similarly, recording of the inhibitory miniature postsynaptic currents showed a significant enhancement in the frequency of these events (SH: 1.19 ± 0.12Hz, PE: 1.82 ± 0.17 Hz, p = 0.005) but no change in the amplitude of mIPSC (SH: 21.10 ± 1.58 pA, PE: 18.11 ± 3.25 pA, p = 0.37). These results suggest that either an increase of the number of excitatory and inhibitory inputs on GC, or a change in the release probability of the existing inputs, or both. To establish whether the increase in mEPSC frequency correlates with an increase of excitatory synapses in the ML of the DG, we performed immunohistochemistry using an antibody targeting the excitatory presynaptic protein vesicular glutamate transporter 1 (VGLUT1), a marker of choice for glutamatergic nerve terminals and glutamatergic synapses (Micheva et al., 2010). Labeling and imaging of brain sections obtained from animals raised in either the SH or the PE condition demonstrates an increase in the density of VGLUT1 puncta in the ML of mice raised in EE. This increase in VGLUT1 puncta density reflects an increase of excitatory synapses formed onto GC dendrites and is in accordance with previous studies (Hosseiny et al., 2015). The observed increase of mIPSC frequency in our experiment suggests that the circuit is adding inhibitory synapses in order to maintain a proper excitatory-inhibitory balance in the DG. While an increase in mIPSC frequency has previously been observed in the DG of mice raised in an EE (He, Ma, Liu, & Yu, 2010; Liu et al., 2012), the cellular origin of the enhanced inhibitory synaptic drive is unclear. We therefore designed a series of experiments to test whether specific inhibitory cell-types are more sensitive to experience-dependent plasticity in the DG.

**Figure 1.**
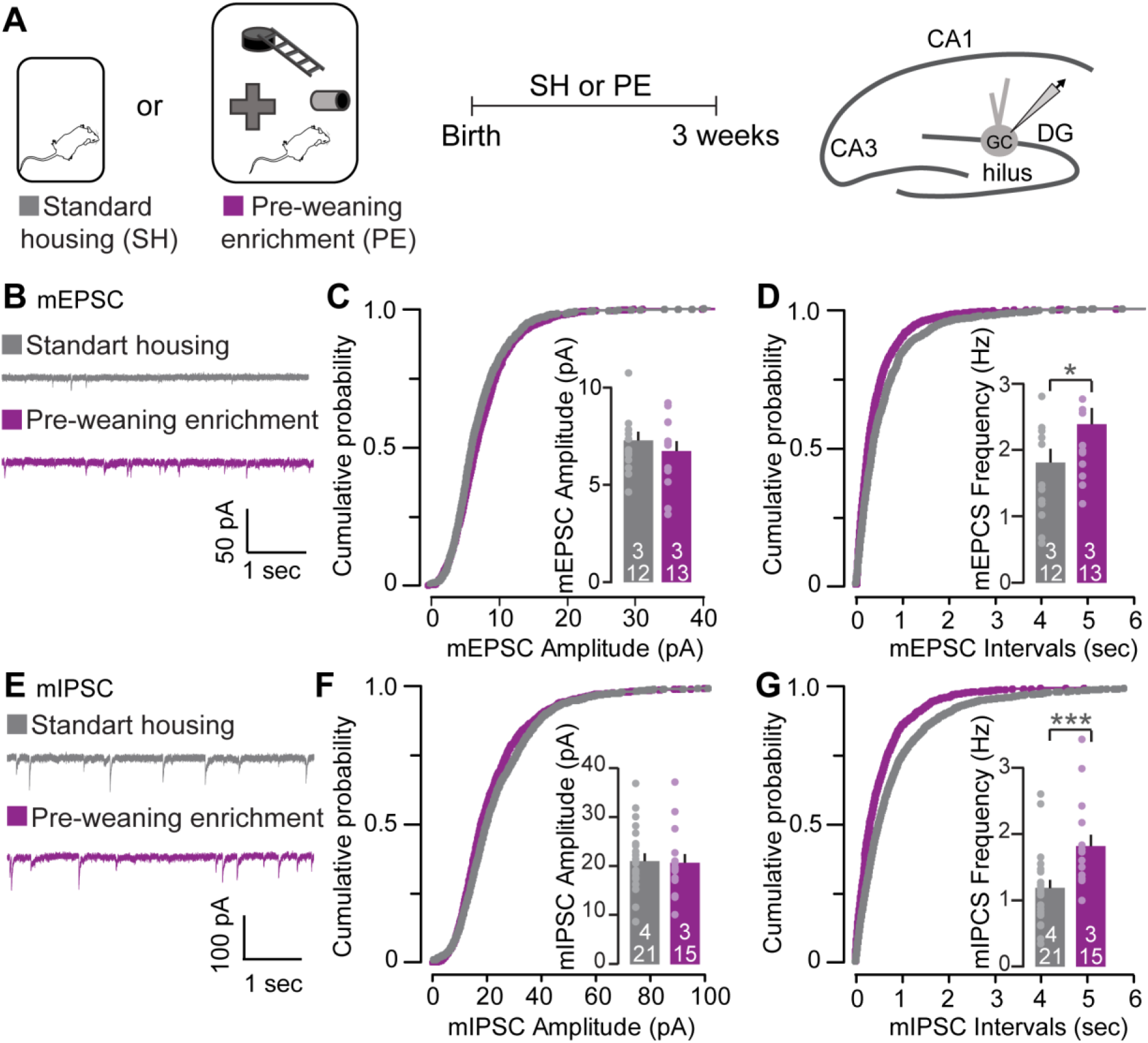
PE increases inhibitory and excitatory drive in the DG. (**A**) Illustration of the standard housing (SH) and pre-weaning enrichment (PE). Mice were raised either in SH or PE for 3 weeks after birth. Schematic illustration of the hippocampus with a representative whole cell patch clamp recording of a granule cells in the DG. (**B**) Representative mEPSC traces recorded from mature GCs (P19-21) in acute brain slice from SH and PE. (**C** and **D**) Cumulative distribution of the inter-event intervals (C) and amplitude (D) of the mEPSC recorded in GCs from SH and PE (P19-P21). Insets: mean frequency and amplitude. *p < 0.05 by two-sample t-test, SH: a/n = 3/12, PE: a/n = 3/13. (**E** to **G**) Same as B to D but for mIPSC. ***p < 0.001 by a two-sample t-test, SH: a/n = 4/21, PE: a/n = 3/15. Data represent mean ± SEM. For each bars, the number of animals and the number of recorded cells used for the quantification is reported (a/n).

### PE increases GC perisomatic inhibition

The dentate gyrus is populated by a diverse population of inhibitory interneurons with distinct morphological and molecular characteristics. These neurons innervate their axons to the specific laminae within the DG and provide unique inhibitory modalities to GCs. For instance, the soma of somatostatin-expressing cells are present in the hilus and contact the apical dendrites of GC in the molecular layer (ML), providing weak and slow dendritic inhibition (Savanthrapadian et al., 2014; Yuan et al., 2017). On the other hand, perisomatic inhibition is controlled by both the CCK+ and PV+ basket-cells that are present in the inner granular cell layer (GCL) and, in accordance with this propitious subcellular targeting, these cell-types exert strong temporal control on GC action potential generation (Bartos & Elgueta, 2012; Hefft & Jonas, 2005) (**Figure 2A**). Using an antibody targeting vesicular GABA transporter (VGAT) present at all inhibitory presynaptic terminals, we were able to quantify the density of inhibitory synapses present in the different layers of the DG (**Figure 2A**). We measured VGAT puncta density in the ML, inner molecular layer (iML), GCL and hilus using 3D puncta quantification. We found a specific increase in VGAT puncta density in ML and GCL (ML: SH: 100 ± 3.89%, PE: 115.61 ± 3.32 %, p = 0.0046; GCL: SH: 100 ± 4.42 %, PE: 141.29 ± 7.59 %, p = 0.00024), but no significant change in the hilus or in iML (hilus: SH: 100 ± 4.03%, PE: 94.90 ± 4.84 %, p = 0.53; iML: PE: 106.85 ± 7.09 %, SH: 100 ± 8.34%, p = 0.55) (**Figure 2C**). This result suggests that the increased inhibitory drive on the GC observed with mIPSC recording is, at least in part, due to the structural plasticity involving an increase of dendritic and somatic inhibition onto GCs. To confirm that an increase in VGAT puncta density reflects the addition of inhibitory synaptic inputs onto the GC, we used a viral approach to express the inhibitory postsynaptic protein, gephyrin fused to GFP in GC. Viral injection of AAVDJ-hSyn-GFP-Gephyrin were done at P1, and the brain was harvested after mice were raised in SH or EE for three weeks (**Figure 2D**). We quantified Gephyrin-GFP clusters on the somatodendritic axis of the GCs across the different laminae (**Figure 2D** and **S2**). In accordance with an increase in VGAT puncta density in the GCL, we found a significant increase of gephyrin-GFP clusters present on GC somas and proximal dendrites (PD) (soma: SH: 34.50 ± 1.68, PE: 44.42 ± 2.72, p = 0.0029; PD: SH: 0.34 ± 0.04, PE: 0.43 ± 0.03, P = 0.039) (**Figure 2D**). The number of Gephyrin-GFP puncta on the GC axon initial segments (AIS) was not affected by EE (SH: 0.39 ± 0.03; PE: 0.39 ± 0.05, p = 0.945), which suggests that synaptic inputs from axo-axonic inhibitory cells such as PV^+^ chandelier cells are not altered by EE. Surprisingly, despite a mild but significant increase of VGAT puncta density in the ML following EE (**Figure 2C**), no increase of Gephyrin-GFP clusters were found on the dendrites of the GC in PE mice (DD: SH: 0.28 ± 0.01, PE: 0.28 ± 0.01, p = 0.762; MD: SH: 0.25 ± 0.02, PE: 0.23 ±0.01, p = 0.422) (**Figure 2D**). Altogether, these results suggest that enhanced spontaneous inhibitory activity on GCs is due to an increase in somatic inhibition on GCs and demonstrate that experience-dependent synaptic plasticity is restricted to specific inhibitory cell-types that target GC soma.

**Figure 2.**
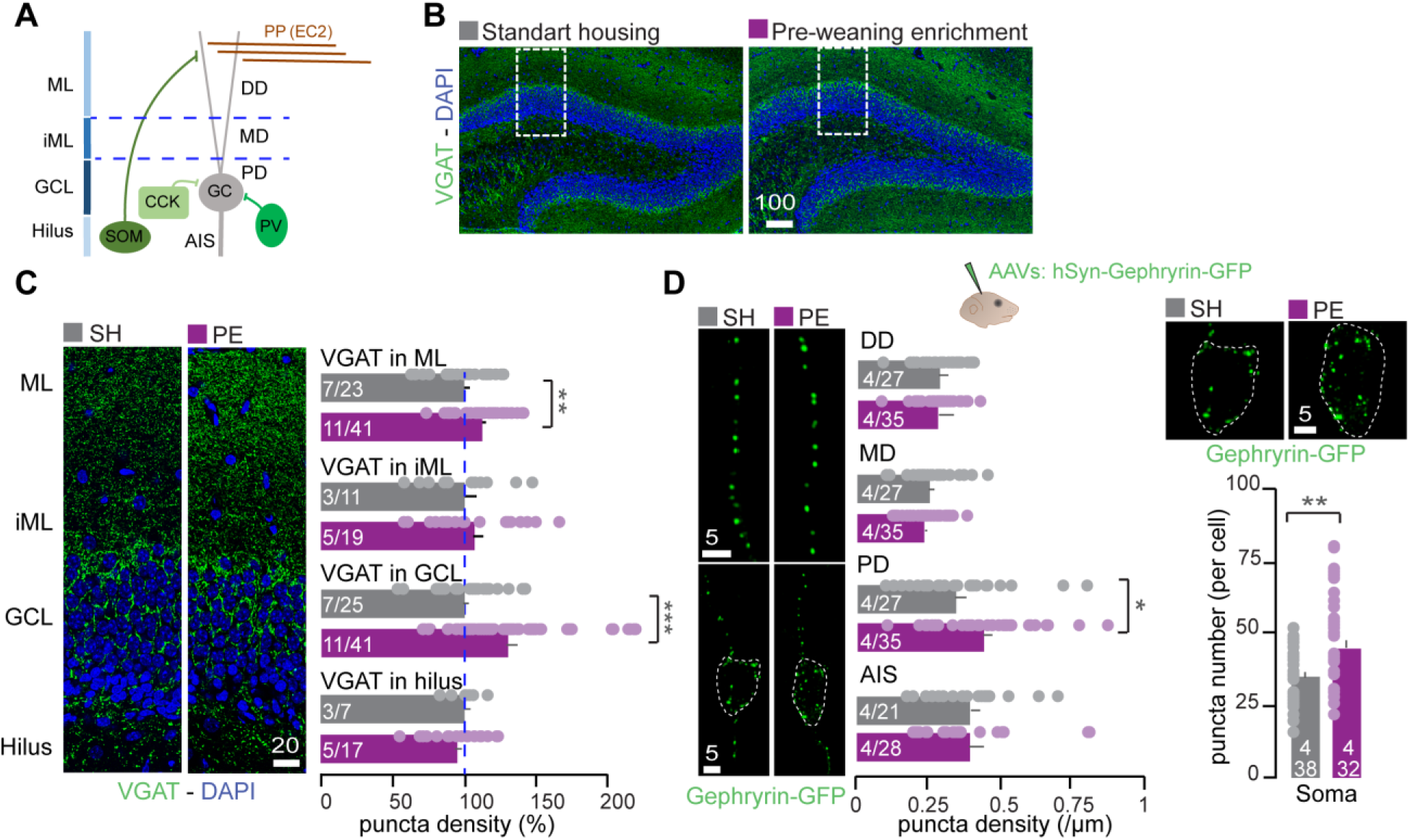
PE induces a perisomatic increase of inhibitory synapses on GC. (**A**) Schematic representation of interneuron connectivity in the DG. GC, granule cell; PP(EC2), the perforant path i.e. projection from layer 2 entorhinal cortex neurons; PV, parvalbumin interneuron; CCK, cholecystokinin interneuron; SOM, somatostatin interneuron. Dendrites are segmented based on their location within the different laminae; distal dendrites (DD) in the molecular layer (ML), medial dendrites (MD) in the inner ML, proximal dendrites (PD) in the GC layer (GCL) and axon initial segment (AIS) in the hilus. (**B**) Confocal images of the DG from SH (left) and PE (right) mice stained with anti-VGAT (green) antibody and DAPI (blue). Box with dashed line indicates the region acquired at high magnification for VGAT puncta density quantification. (**C**) Left, representative images of the VGAT labeling for SH and PE mice. Right, quantification of VGAT puncta density in all laminae of the DG. (**D**) Top; schematic of the adeno-associated virus (AAV) injection in neonatal pups at P1. Left; representative images of gephyrin-GFP fluorescence along the somatodendritic axis of GCs from SH and PE mice. The quantifications of the density of GFP clusters along the segments of the dendrites and the AIS are reported as number of clusters per μm. Dashed lines represent the contour of the cell soma. Right, representative images of GC soma labeled with gephyrin-GFP and quantification of the number of cluster per soma. Data represent mean ± SEM. Statistic by a two-sample t-test; *p<0.05, **p < 0.01 and ***p < 0.001. For each bars, the number of animals and the number of images used for the quantification are reported (a/n). All scale bars are in μm.

### CCK+ but not PV+ somatic inputs are increased by enriched environment

Two types of basket cells form perisomatic inhibitory inputs onto GCs. Despite shared anatomical similarities, they differ in the way they gate GC activity. This is due in part to the differences in their connectivity patterns, intrinsic excitability, and differential expression of receptors and channels that modulate their activity. For instance, Synaptotagmin 2 is enriched in PV+ basket cell terminals, which is critical to sustain high frequency GABA release (Chen, Arai, Satterfield, Young, & Jonas, 2017; Sommeijer & Levelt, 2012). CCK+ basket cells, on the other hand, express Cannabinoid receptor type 1 (CB1R) at their synaptic terminals and are therefore sensitive to cannabinoid retrograde signaling (Katona et al., 1999). Considering their critical roles in information processing, we questioned how each cell types contributes to the increased somatic inhibition caused by EE. To this end, we first labeled the somatic innervation of PV+ and CCK+ axon terminals using antibodies against Syt2 and CB1R respectively (**Figure 3** and **S3**). Brain sections were obtained from 3 week-old mice raised in SH or EE. Then, the puncta density was quantified (**Figure 3A**). We found that while CB1R/VGAT puncta density was increased by 26%, no change in Syt2 puncta density was found (CB1R/VGAT: SH: 100 ± 10.2%, PE: 126.10 ± 5.31%, p = 0.021; Syt2: SH: 100 ± 2.5%, PE: 103.86 ± 2.57%, p = 0.328). The result demonstrates that the density of CCK+ synapses on GC soma is increased by EE while PV+ inputs are unchanged. Next, we asked whether the increase in mIPSC frequency observed in the PE group (**Figure 1D**) could be reversed by specifically blocking spontaneous release at inhibitory CCK+ synapses. For this purpose we used ω-conotoxin GVIa, a N-type voltage-gated calcium channel (VGCC) blocker, specifically blocking mIPSC from CCK+ terminals that express CB1R (Goswami, Bucurenciu, & Jonas, 2012). We observed that the increase in mIPSC frequency in EE condition was almost completely blocked by bath application of ω-conotoxin GVIa (1 μM) (SH: 1.07 ± 0.11Hz, SH with blocker: 0.99 ± 0.18Hz PE: 1.59 ± 0.13 Hz, PE with blocker: 1.07 ± 0.13Hz, F(3,30) = 4.495, p = 0.01). These strongly suggest that the increased somatic inhibition induced by PE is due to the selective addition of CCK+ inhibitory somatic inputs onto GCs.

**Figure 3.**
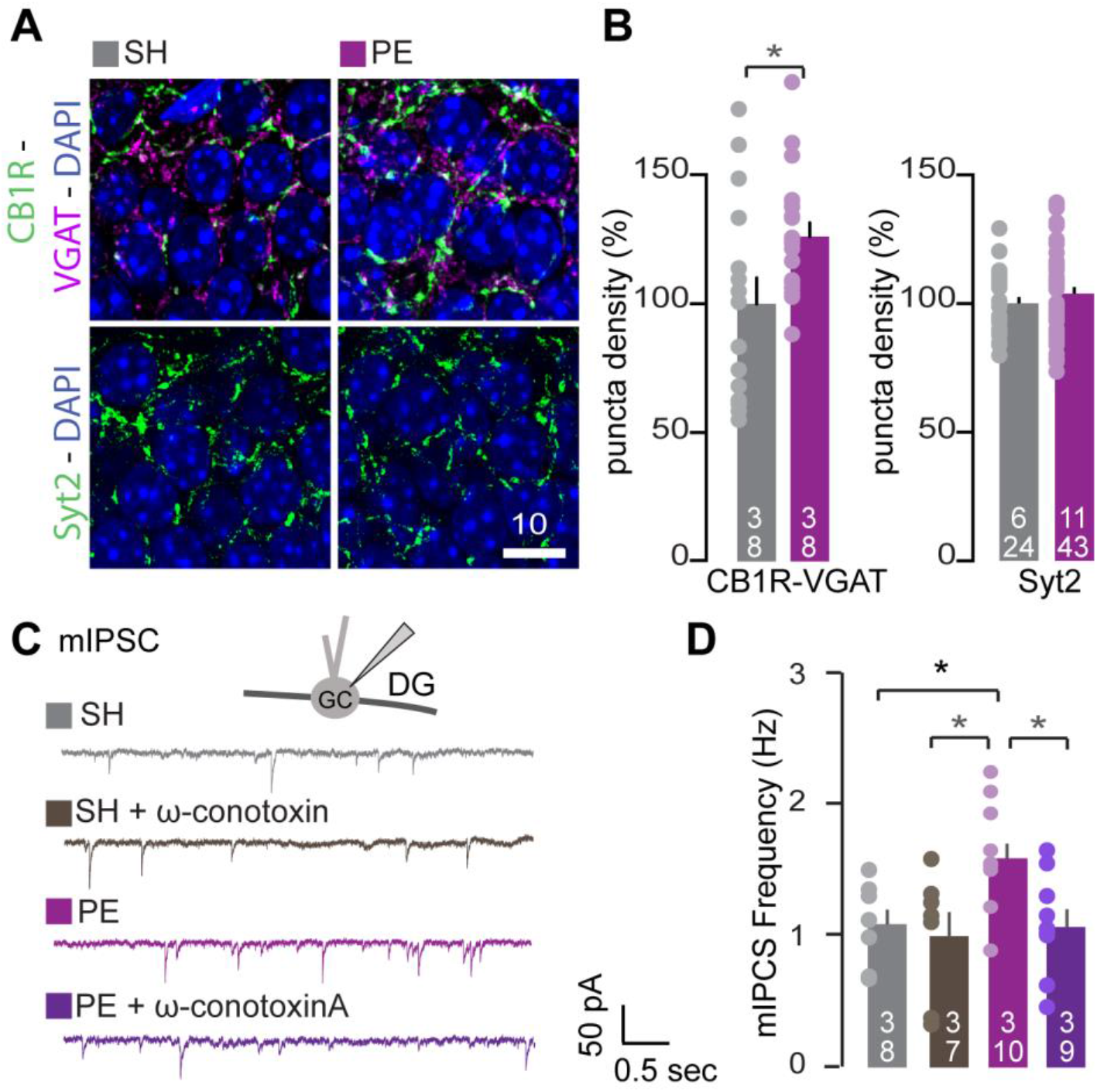
PE enhances CCK+ somatic synapses in the DG. (**A**) Representative images of the GC layer stained with either antibodies against VGAT (purple) and CB1R (green), or Syt2 (green) from brain sections of SH and PE mice. Scale bar = 10μm. (**B**) Corresponding quantification of CCK + basket cell synapses (overlap of VGAT and CB1R) and PV+ synapses (Syt2) in the GC layer. Statistic by a two-sample t-test; *p<0.05. (**C**) Representative mIPSC traces recorded from mouse GCs (P19-21) in the absence or presence of 1 μM extracellular ω-conotoxin GVIA from SH and PE mice. (**D**) Summary graphs of the average frequency. Statistical comparisons were performed with a one way analysis of variance (ANOVA; *p<0.05) followed by Tukey HSD post-hock. Data represent mean ± SEM. For each bars, the number of animals and the number of images or recorded cells used for the quantification are reported (a/n).

### Bidirectional plasticity of somatic CCK+ innervation is controlled by the activity of the perforant path

In the DG, the fibers of the perforant path (PP) originate from the EC2 long-range projection neurons. Their axons carry multimodal sensory information and form the major excitatory inputs in the circuit (Knierim, Neunuebel, & Deshmukh, 2014; Witter & Moser, 2006). Previous work have demonstrated that the activity of EC2 afferents has an instructional role in DG maturation (Donato, Jacobsen, Moser, & Moser, 2017; Pieraut et al., 2014). We therefore reasoned that activity from these afferents could also regulate the plasticity of CCK+ interneurons. To test this hypothesis, we manipulated activity of EC2 neurons using viral-mediated expression of the Designer Receptors Exclusively Activated by Designer Drugs (DREADD) receptor hM4D(Gi). First, we tested whether expression of hM4D(Gi) and its activation with clozapine-N-oxide (CNO) was sufficient to suppress the experience-dependent expression of an immediate early gene, cFOS in the DG. Naïve mice previously injected with AAV-hSyn-hM4D(Gi)-mCherry in the EC at P1 were placed in an EE cage at 3 weeks of age. The expression of cFOS was assessed 90mins later using immunostaining. Quantification of cFOS+ cells in the DG demonstrated that CNO injections repressed the experience-dependent expression of cFOS in the GCL from the mice expressing hM4D in the EC (**Figure S4**). Having demonstrated that our approach enabled the manipulation of activity of the EC projection neurons, we then chronically inhibited these EC afferents in naïve animals raised in SH condition with twice daily intraperitoneal injections of CNO from P15 to P21 (**Figure 4A**). We then stained brain sections with the inhibitory synaptic markers to visualize CCK+ and PV+ inhibitory synapses respectively (VGAT and CB1R in one hand or Syt2 alone) (**Figure 4B**). Quantification demonstrates that chronic inhibition of the PP led to a reduction of somatic CCK+ inhibitory synapses on the GC (SH-control: 100 ± 7.10%, SH-silenced: 59.678 ± 5.39%, p = 0.0005). No change in Syt2 puncta density in the GCL was found in the same experimental condition (SH-control: 100 ± 5.760%, SH-silenced: 97.370 ± 5.190% p = 0.752) (**Figure 4C**). Thus, the activity of EC afferents during the third week of postnatal development is critical for the maturation of inhibitory synapses formed by CCK+ basket cells, but not the ones formed by PV+ cells. In light of this finding, we reasoned that the activity of EC2 afferents could be the driving signal regulating inhibitory plasticity in the opposite direction (i.e. addition of CCK+ synapses) as observed in mice raised in EE (**Figure 3**). We therefore tested whether hM4D-dependent inhibition of the EC2 projections would prevent the increase of CCK+ somatic synapses observed in the PE group. In accordance with our previous experiments, PE induced an increase of VGAT/CB1R puncta overlap in the GCL, but this increase did not occur when the EC2 afferents were chronically inhibited with the hM4D chemogenetic approach. Importantly, the density of VGAT/CB1R measured in PE mice not expressing hM4D and injected with CNO was similar to that of control PE mice (PE-hM4D-CNO: 102.76 ± 6.67%, PE-hM4D-NS: 144.90 ± 7.69%, SH-hM4D-NS: 100 ± 7.56%, p=0.0026 for PE-hM4D-CNO and PE-hM4D-NS, p < 0.0001 for SH-hM4D-NS and PE-hM4D-NS; F(3 69) = 10.313, p < 0.0001) (**Figure 4E**). These demonstrate that activity of EC2 afferents is critical for the experience-dependent plasticity of CCK+ somatic inhibition.

**Figure 4.**
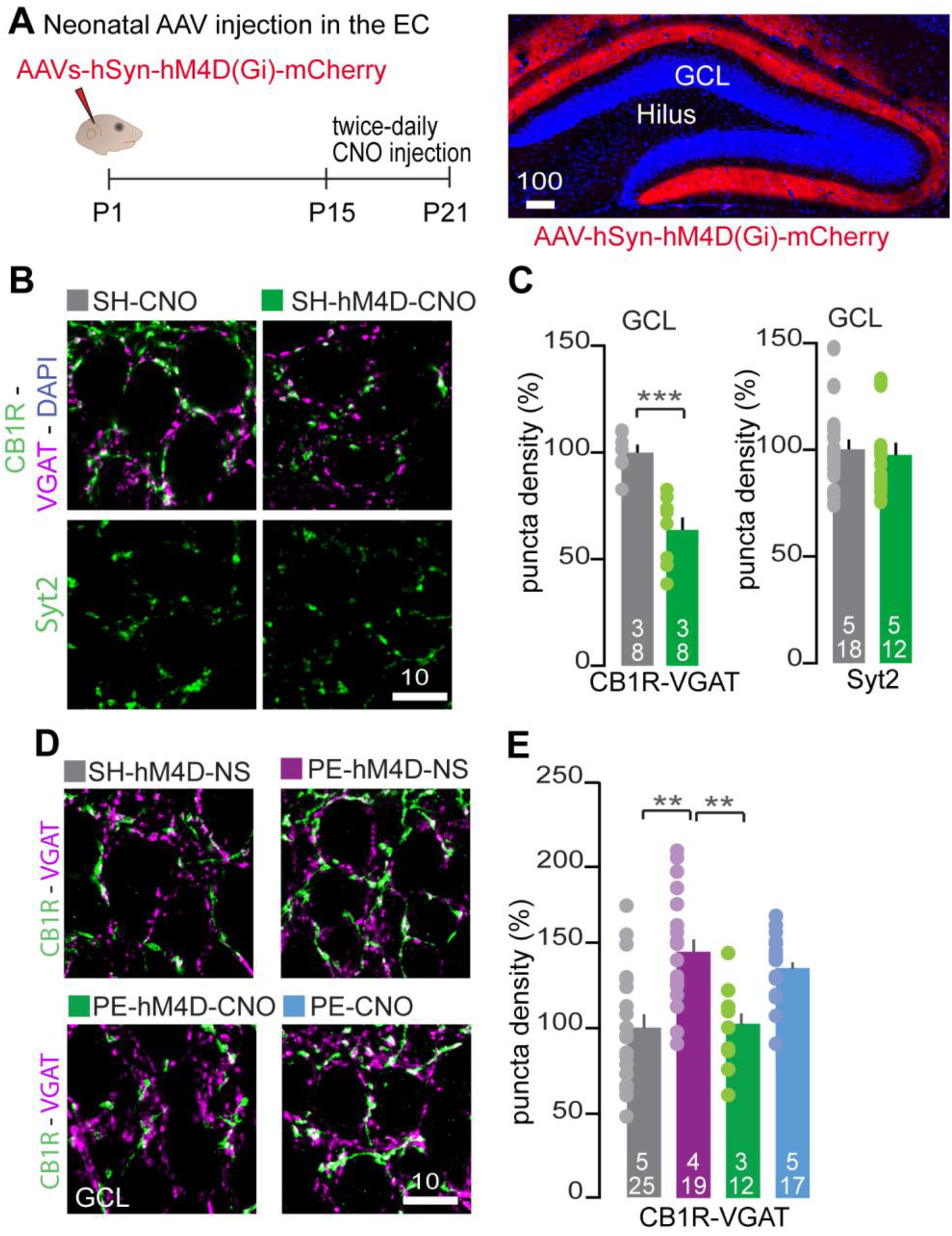
Experience-dependent remodeling of CCK+ basket cell inputs is controlled by EC2 neuron activity. (**A**) Left, experimental paradigm for chemogenetic inhibition of excitatory afferents from EC2 in the DG circuit. Right, representative image of the perforant path fibers in the DG (afferents from the EC2) labelled with mCherry (Red) following neonatal injection of AAV-hSyn-hM4D(Gi)-mCherry in the EC. (**B**) Representative images of the GC layer stained with antibodies against either, VGAT (purple) and CB1R (green), or Syt2 (green) from brain sections of SH mice injected or not with hM4D expressing virus and treated with CNO (SH-hM4D-CNO and SH-CNO respectively). (**C**) Corresponding quantification of CCK+ synapses (left) and PV+ synapses (Syt2) (right) in the GC layer. Graph bars represent mean ± SEM. All data were normalized to SH-CNO control. Statistical comparisons were performed with a two-sample t-test (***: p<0.001). (**D**) Same as for (A) but chronic inhibition of the EC2 afferents was done in PE mice (PE-hM4D-CNO) and compared to values obtained from SH and PE mice also injected with the virus but who received NS treatment (SH-hM4D-NS and PE-hM4D-NS respectively). An additional PE control group with no virus received the CNO injection (PE-CNO). (**E**) Corresponding quantification of CCK+ synapses in the GC layer of brain sections for each condition. Statistical comparisons were performed with an ANOVA (**: p<0.01) followed by Tukey HSD post-hock test. For each bars, the number of animals and the number of images used for the quantification are reported (a/n). All scale bar are in μm.

### Pre-weaning enrichment has limited effect on pattern separation in the DG

In the DG, the connectivity of the GABAergic network is uniquely suited to support pattern separation (Bel et al., 2015; Espinoza et al., 2018; C.-T. T. Lee et al., 2016). In light of the structural remodeling induced by EE observed in our study, we questioned whether these changes could impact the transformation of the entorhinal inputs in the circuit. To test our hypothesis, we decided to use an electrophysiological approach recently established by Madar and colleagues, enabling us to assess pattern separation in the DG (Madar, Ewell, & Jones, 2019a). This protocol enables to compare the similarity of the incoming signal (input spike trains) delivered to the DG with the similarity of the recorded output spike trains. We used slice physiology to stimulate afferents inputs with 5 distinct 10Hz input spike trains and to record the output spike trains recorded on single GCs simultaneously. The similarity of the inputs and outputs can be calculated using multiple metrics, each of which inform about computational transformation performed by the DG (**Figure 5A**) (Madar et al., 2019a). The spiking activity of the cells was first computed to calculate the output correlation (R_output_) using pairwise Pearson’s correlation coefficient (R). We found that in both SH and PE mice, R_output_ was lower that the R_input_, supporting a decorrelation of the incoming signal by DG. This is in accordance with pattern separation operated in this circuit. When comparing the mean R_output_ obtained from SH and PE mice, we found a statistically significant decrease but the difference between the two groups was very small (mean SH-R_output_ = 0.27, mean PE-R_output_ mean = 0.25, p = 0.029). Moreover, while statistically different, correlation coefficients of the R_output_ with the R_input_ are very close between the two groups (**Figure 5B**). Because pattern separation can be considered as having the inputs’ pattern being orthogonalized, we also computed the normalized dot product (NDP, cosine of the angle between two vectors) of both input and output patterns. This allowed us to assess the extent to which the signal, viewed as vectors of spike-counts, is transformed by the DG (i.e. “orthogonalized”). We found no statistical differences between SH and PE mice for the NDP metric (mean SH-NDPoutput =0.30, mean PE-NDPoutput =0.28, p = 0.14) (**Figure 5B**). Lastly, we computed the data to compare the scaling factor value (SF) of the input and output spike trains. This metric enabled us to assess if the number of spikes per “bin” is scaled up or down. Our analysis demonstrated a small but statistically significant change of the SFoutput values in the PE group (mean SH-SFoutput =0.83, mean PE-SFoutput =0.88, p = 0.000) (**Figure 5B**). While SF, but not R or NDP, is dependent on the firing rate (Madar et al., 2019b), we wanted to assess if the firing property of the outputs were different among the two groups. We first computed the data to compare the firing rate (FR) and burstiness (pBurst) of the recorded neurons. Both pBurst, i.e. the probability of firing a small burst of AP following a single stimulus, and firing rate were identical in SH and PE groups (**Figure 5D**). To further explore a putative change in spike train features, we calculated dispersion of the spike train firing rate, compactness, and occupancy (see material and method section for more details). We found no difference in these metrics, suggesting that in response to the same input pattern, the output spike trains are highly similar in term of pattern of spike distribution and firing structures. While small but statistically significant differences were found between the two groups for SF and R metrics, we argue that to a large extent, pattern separation at the cellular level is not affected by housing condition.

**Figure 5.**
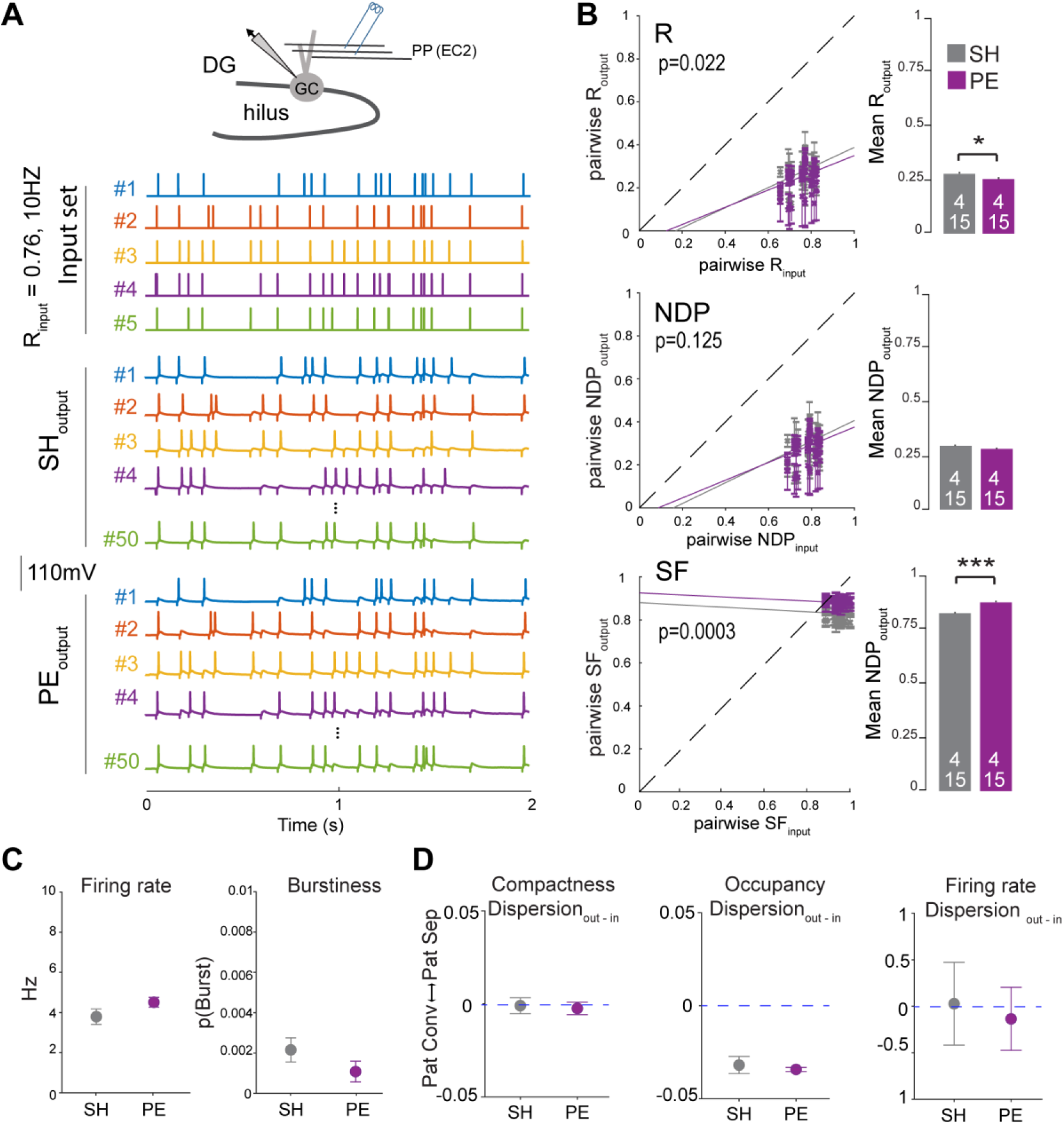
PE housing has limited effect on pattern separation in the DG. (**A**) Top, cartoon depicting whole-cell recording of the GCs during simultaneous stimulation of the PP. Bottom, representative traces obtained in current-clamp mode in response to the stimulation with 5 input spike trains. To assess the level of pattern separation operated by single recorded GCs, the similarity of the inputs is compared to the similarity of the outputs using different comparative metrics. (**B**) Representative graphs of pattern separation with pairwise output similarity versus the pairwise input similarity measured by the three different metrics, including R, NDP and SF and using 10 ms bins. Data points below and above the dashed line correspond to pattern separation and pattern convergence, respectively. Solid lines represent the linear fits for the ANCOVA. Statistical comparisons were performed with an ANCOVA (the aoctool function in Matlab) and a two-sample t-test (*: p<0.05 and ***: p<0.001). (**C**) A summary for FR and p(Burst) (number of animals = 4 for both group, total 15 recordings for each group). Statistical comparisons were performed with a two-samples t-test for firing rate (t = 1.559, p = 0.13) and p(Burst) (t = 1.3627, p = 0.184). (**D**) Compactness, Occupancy or FR codes were measured for each recording sets. Statistical comparisons were performed with 2 sample t-test for binwise Compactness of output (t = 1.04, p = 0.307), the variations of Occupancy (t = 1.924, p = 0.073) and the variations in FR or spike trains (t = 1.129, p = 0.269). For each bars, the number of animals and the number of recorded cells used for the quantification are reported (a/n).

## DISCUSSION

Pre-weaning enrichment enables to assess how enhanced experiences driven by increased exploratory activity, sensory stimuli, and social interactions can affect the maturation of the synaptic network during postnatal development. Consistent with previous work, we found that this housing paradigm increases excitatory and inhibitory drive in the DG (**Figure 1**) (see also (Hosseiny et al., 2015; Liu et al., 2012)). These findings raise a critical question: what is the cellular origin of this plasticity?

In this study, we characterized a new form of plasticity that affects the formation of CCK+ inputs onto the soma of GC in the DG. Using a combination of enrichment housing and chemogenetic approaches, we demonstrate that the maturation of CCK+ basket cell connectivity is strongly governed by experience and requires the activation of the afferents from the EC2. By combining immunohistochemistry with cell-type specific markers and genetic labeling with gephyrin-GFP, we first demonstrate that EE leads to an increase in inhibitory synapses around the soma of the GC (**Figure 2**). Immuno-labeling of synaptic markers specific to PV+ and CCK+ basket cells were instrumental to distinguish the two well characterized basket cell-types in the DG and reveals that terminals from CCK+ but not PV+ basket cells are increased by environmental enrichment. This conclusion is strongly supported by the fact that the experience-dependent increase of mIPSC frequency in PE mice is blocked by ω-conotoxin GVIa, a N-type VGCC blocker previously shown to block mIPSC originating from CCK+ terminals (Goswami et al., 2012). Together, these experiments support the conclusion that in response to PE, perisomatic addition of CCK+ basket cells terminals increase GC inhibition.

It is possible that the limitation of our imaging and electrophysiological approaches prevents us from finding the implication of other IN populations in the remodeling of the inhibitory synaptic network in PE mice. For instance, recording of mIPSC on cell soma skews the sampling of synaptic events in favor of those originating from the perisomatic and proximal dendritic domains and at the disadvantage of those originating from the distal part of the dendrites (Soltesz, Smetters, & Mody, 1995). Thus, recording of mIPSC may lack the sensitivity needed to reveal the remodeling of dendritic targeting INs such as the SST+ hilar perforant path (HIPP) cells. This could explain why bath application of ω-conotoxin fully blocked the increase of mIPSC frequency despite the fact that we found an accumulation of VGAT puncta in the ML after PE. However, the change in VGAT puncta density in ML was modest and did not correlate with an increase of Gephyrin-GFP clusters in distal dendrites. A change in VGAT puncta density was also observed following manipulation of PP activity with hM4D (**Figure S5**). It is possible that the change in VGAT puncta density in the ML reflects the addition of inhibitory synapses on inhibitory dendrites and not on excitatory GC dendrites. This could explain the discrepancy between VGAT and Gephyrin-GFP analysis. Another hypothesis is that axon terminals from INs are added but the maturation of the postsynaptic counterpart is delayed. The later hypothesis is consistent with in vitro work showing that VGAT clustering precedes that of Gephyrin (Dobie & Craig, 2011; Frias et al., 2019). Future work will be needed to distinguish between these two non-exclusive hypotheses. Nevertheless, based on the evidence presented in this study, CCK+ basket cells play a central role in adjusting inhibition on GC through addition of perisomatic CB1R+ inputs.

The specific recruitment of CCK+ basket cells’ plasticity in response to experiences underscore the existence of molecular mechanisms induced by activity and controlling the formation of somatic inputs on GCs. This is further supported by our chemogenetic silencing of EC afferents with hM4D (**Figure 4**). What are the downstream effectors triggered by EC2 neurons activity and leading to CCK+ plasticity? Using PE paradigm, previous studies have shown that housing condition affects the release of neurotrophins (NTs) such as BDNF in juvenile mice (Liu et al., 2012; Novkovic, Mittmann, & Manahan-vaughan, 2015). It is therefore possible that the increase in synaptic connectivity observed in PE mice is due to a change in release of NTs. BDNF has been widely studied and many experiments demonstrate its role in the control of excitatory as well as inhibitory synapse formation (Fiorentino et al., 2009; Sakata et al., 2009; Sallert et al., 2009). Here, we have demonstrated that manipulating activity of the perforant path is able to prevent experience-dependent remodeling of the CCK+ inputs on GC (**Figure 4**). This bidirectional control of CCK+ innervation by PP activity may be mediated by the release of NTs in the circuit. To support such a hypothesis, it would be interesting to assess the extent to which activity of the entorhinal inputs regulate the release of NTs in the DG and determine which cell type is involved in such release. Activity-driven expression of transcription factor is also a very likely scenario in the control of CCK+ plasticity. Notably, it is possible that activity of EC afferents directly activates expression of immediate early genes such as Neuronal PAS Domain Protein 4 (NPAS4) that has a role in experience-dependent remodeling of the inhibitory network in the CA1 (Bloodgood, Sharma, Browne, Trepman, & Greenberg, 2013). In fact, in this circuit, enriched environment led to an increase in somatic inputs from CCK+ cells onto the pyramidal cells in response to NPAS4 expression (Hartzell et al., 2018). NPAS4 in GC may activate specific transcriptional programs and induce the release of retrograde signaling molecules such as NTs. BDNF was actually linked to NPAS4 mediated remodeling of the inhibitory network in the CA1 (Bloodgood et al., 2013). However, it is surprising that the chronic blockade of EC afferent activity with hM4D induces the remodeling of CCK+ but not PV+ terminals. In a previous work, silencing of these fibers with tetanus toxins light chain considerably decreased the density of PV+ inputs on GC (Pieraut et al., 2014). However, while expression of tetanus toxin totally abolishes the release of neurotransmitter, hM4D hyperpolarizes cells and thus may have a distinct effect. Also, expression of tetanus toxin can block the release of BDNF (Shimojo et al., 2015) and may affect PV+ innervation through a distinct mechanism. Another interesting hypothesis is the timing of the manipulation of EC activity. The silencing with tetanus was taking place at P3-4 postnatally, whereas CNO injections started at P15 in our study. Assessing how silencing of the afferents affects the assembly of the inhibitory network at different developmental time windows could reveal the existence of a hierarchy in the maturation of distinct classes of IN. Interestingly, the maturation of the different subfields along the entorhino-hippocampal circuit is controlled by excitatory activity that spreads in a hierarchical bottom-up manner along this linear circuit (Donato et al., 2017). This study, along with the work presented here, supports a model in which activities from upstream brain regions have an instructional role in the maturation of microcircuits along the entorhino-hippocampal circuit. This spatial and temporal maturation of the circuit recapitulates the flow of excitation through the circuit and must have a critical role in establishing the connectivity rules required for the proper computation within each microcircuit.

Recruitment of GC during spatial exploration and learning tasks is very sparse (Chawla et al., 2005; Deng et al., 2013; Hainmueller & Bartos, 2018; Leutgeb, Leutgeb, Moser, & Moser, 2007). Anatomical and electrophysiological studies demonstrate that the unique properties of the inhibitory network in the DG is well suited to support sparsification and decorrelation (Acsády, Kamondi, Sík, Freund, & Buzsáki, 1998; Acsády, Katona, Martínez-Guijarro, Buzsáki, & Freund, 2000; Bel et al., 2015; Espinoza et al., 2018; C.-T. T. Lee et al., 2016; Madar et al., 2019b; Sambandan, Sauer, Vida, & Bartos, 2010). Nonetheless, the specific role of distinct classes of IN in DG computation is not well established. Considering the strong reorganization of the inhibitory network induced by EE, we attempted to ascertain whether the decorrelation of cortical inputs in the DG was different in SH and PE mice. By testing different similarity metrics, we find that decorrelation of the inputs is strongly preserved at the cellular level when the inputs are set at 10Hz. While two similarity metrics, R and SF, show statistical differences, the observed effect sizes are small. We therefore conclude that the GC performs pattern separation through a high degree of othogonalization (R and NDP) and low levels of scaling (SF) independent of the housing condition (**Figure 5**). Owing to the fact that frequency of the inputs has a strong influence on GC function (Bel et al., 2015; Madar et al., 2019b), more work would help to determine whether the change in CCK+ innervation can affect pattern separation at specific input frequencies. It is also important to note that at the population level, pattern separation could be affected by a change in somatic inhibition. CCK+ cells have slow and asynchronous inhibition strongly controlled by neuromodulators (Bartos & Elgueta, 2012; Földy, Lee, Szabadics, Neu, & Soltesz, 2007; Hefft & Jonas, 2005; S. H. Lee & Soltesz, 2011). Activation of CB1R at CCK+ basket cell terminals induces depolarization-induced suppression of inhibition (DSI), a process by which repetitive activation of GC leads to disinhibition from CCK+ cells. GC disinhibition through this mechanism can favor their firing and support a “winner take all” situation, increasing activity of the sparse GCs already active. Therefore, a change in somatic inhibition from CCK+ basket cell terminals expressing CB1R can favor sparsification of the entorhinal signal and increase the signal to noise ratio. DSI-dependent GC disinhibition may also favor plasticity at excitatory inputs of synaptically connected mossy cells and CA3 pyramidal cells (Rebola, Carta, Lanore, Blanchet, & Mulle, 2011). Such mechanisms would promote associative pairings critical for learning and memory formation. Taking advantage of cell-type specific genetic targeting in combination with optogenetic or chemogenetic manipulation of cell activity, future work will be needed to dissect the distinct, yet complementary role of the varied inhibitory cell-types in DG computation (C.-T. T. Lee et al., 2016).

In light of the findings demonstrating the beneficial role of PE in mouse models of neurodevelopmental disorders such as Down syndrome and Rett syndrome (Begenisic, Sansevero, Baroncelli, Cioni, & Sale, 2015; Lonetti et al., 2010), a better understanding of the cellular and molecular changes promoted by early experiences is desirable. Moreover, considering that disruption of CCK+ cells connectivity impact the generation of theta oscillations and memory formation in adult (del Pino et al., 2017), establishing the mechanisms underlying the synaptic integration of this cell-type is also needed. In the present study, we demonstrate that among the varied inhibitory synapses in the DG, somatic CCK+ innervation is adjusted in response to cortical activity. This structural plasticity is engaged during maturation of the DG in an experience-dependent manner. Future work should focus on understanding the signaling mechanisms regulating CCK+ plasticity and exploring the role of this cell type in DG computation.

## Materials and methods

### Animal husbandry and EE raising

All animal procedures were approved by the university of Nevada Reno Institutional Animal Care and Use Committee, which were in accordance with federal guidelines. All experiments were performed on C57BL/6 mice, and both female and male mice were used. All mice were exposed to a 12 h light/12 h dark cycle with food and water provided ad libitum. For pre-weaning enrichment, female mice at embryonic days 16–19 (E16 –19) were randomly assigned to enrichment housing 4 –7 d before delivery with another female companion. The enrichment environment (EE) cage consists of a large Plexiglas laboratory cage (60 × 45 × 20 cm) containing objects of various shapes, colors and textures, including plastic house, tunnels, wood blocks and a running wheel. The different objects and position were rearranged every other day to maximize novelty. All pups were raised in EE cages from birth to juvenile, while the pups that were referred to standard housing were placed with their female breeder in standard control cages. All experiments were performed on animals between postnatal days 19–21 before weaning.

### Acute slice preparation

Transverse hippocampal slices were prepared from C57BL/6L (P19–21). Animals were anesthetized with isoflurane and decapitated. The cerebral hemispheres were removed and bathed for one minute in a cold ice slushy sucrose-based dissection buffer containing (in mM): 87 NaCl, 25 NaHCO3, 1.25 Na2HPO4, 2.5 KCl, 7 MgCl2, 10 glucose, 0.5 CaCl2, 1.3 ascorbic acid, 75 sucrose and equilibrated with 95% O2/5% CO2. Horizontal slices (300 μm thickness) were cut with a VF 300-0Z microtome (Precisionary Instruments) and transferred to a recovery chamber with modified artificial cerebrospinal fluid (ACSF) consisting of (in mM): 92 NaCl, 30 NaHCO3, 1.2 Na2HPO4, 2.5 KCl, 0.5 CaCl2, 10 MgSO4, 25 glucose, 20 HEPES, 5 ascorbic acid, 2 Thiourea and 3 sodium pyruvic, oxygenized with 95% O2/5% CO2. Slices were recovered for 30 min at 32-34 ° C and moved to room temperature for recovering another 30 min before recording, and then maintained at room temperature for the duration of the experiment (4–6 hr).

### Electrophysiology and pharmacology

All recordings were performed in the outer GC layer with a SutterPatch double IPA amplifier (Sutter instrument). Signals were acquired using SutterPatch software (Sutter instrument), filtered at 5kHZ and sampled at 10 kHz. During the recording, slices were maintained in oxygenized standard ACSF, containing (in mM): 124 NaCl, 24 NaHCO3, 1.2 Na2HPO4, 2.5 KCl, 2 CaCl2, 2 MgSO4, 12.5 glucose, 5 HEPES. For miniature inhibitory postsynaptic current (mIPSC) recordings, neurons were clamped at – 70mV, using glass pipettes (resistance 3–5 mΩ) filled with a high chloride internal solution containing the following (in mM): 135 CsCl, 4 MgCl2, 0.1 EGTA, 10 HEPES, 10 Na2-phosphocreatine, 2 Mg-ATP, 0.3 Na-GTP, and 5 QX-314 (pH 7.4, adjusted with KOH to 290-295 mOsm). D-AP5 (Tocris) at 50 μM, CNQX (Torcis) at 10 μM and Tetrodotoxin (TTX)(Enzo) at 1 μM were added to ACSF to block NMDA, AMPA and Na^+^ currents, respectively. For miniature excitatory postsynaptic current (mEPSC) recordings, the GCs were patched with the pipette filled with an internal solution containing (in mM): 122.5 Gluconic acid, 10 CsCl, 122.5 CsOH, 5 NaCl, 1.5 MgCl2, 5 HEPES, 1 EGTA, 10 Na2-phosphocreatine, 3 Mg-ATP, 0.3 Na-GTP, and 5 QX-314 (pH 7.3, adjusted with CsOH to 290-295 mOsm). Neurons were held at –70 mV in the presence of 50 μM picrotoxin (Torcis) to block GABA synaptic currents, and 1 μM TTX. Neurotransmitter release from CCK inhibitory interneurons was pharmacologically blocked by addition of ω-conotoxin GVIa (1 μM) (Sigma and Alomone Labs).

### Virus production and injection

pAAV-hSyn-hM4D(Gi)-mCherry was a gift from Bryan Roth (Addgene viral prep #50475-AAV2; http://n2t.net/addgene: 50475; RRID:Addgene_50475). The virus consist of a human truncated promoter driving the expression of hM4D(Gi) fused to its C-terminal to mCherry (Krashes et al., 2011). The titer of the virus was 5 × 10^12^ viral genomic particles/ml. AAV-hSyn-GFP-Gephyrin were generated with a capsid DJ and purified in our laboratory by a protocol described in (Mcclure, Cole, Wulff, Klugmann, & Murray, 2011). The human Synapsin promoter drives the expression of GFP fused to the N-terminal of gephyrin. The viral titers was 2 x 10^12^ GC/ml, as determined by quantitative PCR. For neonatal in vivo infections, pups were anesthetized on ice and injected with 0.5μl of the viral stock using a glass micropipette (10μm tip diameter). Pups were then warmed up for 5 min on a heating pad and returned to home cages until experiments. The viruses encoding GFP-Gephyrin were delivered into the cerebral lateral ventricles to target GC in the DG (Pieraut et al., 2014). Using the stereotaxic manipulator, the tip of the pipette was placed above lambda to set the zero for both X and Y coordinates. Then, the stereotaxic arms were moved to (X, Y) = (1.0, 1.5) mm for P1 pups. The pipette was inserted into the brain at the depth of 1.5 mm from the skull. For infection of pyramidal neurons with hM4D-mCherry in the entorhinal cortex, pups were laid on one side and the pipette was inserted at 1 mm depth near the bottom of each ear (Pieraut et al., 2014). In this case, expression was restricted to superficial layers of the medial EC. The virus targeting specificity was confirmed in every experiment by imaging mCherry reporter.

### Manipulation of perforant path with DREADD

To manipulate perforant path fibers in the DG, juvenile mice were previously injected with pAAV-hSyn-hM4D(Gi)-mCherry. From P15 to P21, mice received twice daily (9:00 am and 7:00pm) intraperitoneal (IP) injection of either saline (control) or 2.5 mg/kg Clozapine-N-oxide (CNO) solution. Following injection, the juvenile mice were put back with their female breeder. Body weight were monitored every day.

### Immunohistochemistry

Mice were anesthetized with isoflurane and perfused with ice-cold solution containing 4% paraformaldehyde. Brains were post-fixed overnight in 1% paraformaldehyde and sliced in PBS using a vibratome. The 90 mm thick brain sections were blocked in 4% BSA and 0.2% Triton X-100 in PBS at room temperature for 2 h. Sections were then incubated with primary antibodies overnight at 4°C in blocking solution, washed three times in PBS, and incubated in species-matched fluorescently labeled secondary antibodies, and washed 3 times again. Slices were mounted with SlowFade^™^ gold antifade reagent (Invitrogen #S36936) for analysis. Primary antibodies used: rabbit anti-VGlut1 (1:1000; Synaptic System, #135302), mouse anti-VGAT (1:1000; Synaptic System, #131011), rabbit-CB1R (1:1000; Synaptic System, #258003), mouse anti-Synaptotagmin 2 (1:400; DSHB, #AB-23626), Rabbit-cFos (1:1000; Synaptic System, #226003), rabbit-NPAS4 (1:1000; Activity signaling, #AB18A). Fluorescent secondary antibodies were purchased from Invitrogen (A32728, A11008, A11031, A31573).

### Imaging and quantification

Images were collected at 8-bit depth and 1024 × 1024 pixel resolution and 400Hz under an inverted Leica TCS-SP8 confocal microscope. For synaptic bouton and cluster analysis, images were acquired with a 40×, 1.3 oil objective and zoom factor at 3. For cFos density analysis, images were acquired with a 20 x, 0.75 oil immersion objective. All our images were acquired from at least 3 to 4 hemispheres per mice and images were taken in the dorsal regions of the hippocampus. Image analysis was performed blind to experimental conditions using Imaris 9.5 (Bitplane). 3D isosurfaces were created for each object, and automatic spot detection algorithms were implemented for synaptic and axon puncta detection based on 3D colocalization. For gephyrin-GFP cluster counting, dendrite tracing was performed manually by Filament tools in Imaris. Taking advantage of anatomical cues in combination with labeling of the PP with tdTomato (following viral injection to target EC2 neurons), clusters were counted within distinct dendritic segments present in DG laminae (**Figure S2**). Plotted data represented numbers of puncta per μm. For puncta counting and analysis (**Fig. 2, 3 and 4**), the data was normalized to values obtained from control SH examined in parallel. This normalization was used to account for potential variability in synapse density across independent experiments.

### Pattern separation

To study pattern separation, we used a method developed by Madar and colleagues to quantify the transformation of the incoming signal in the DG by individual GC (Madar et al., 2019a). Briefly, horizontal brain sections containing the hippocampus were prepared from SH and PE mice age P19 to P21. Sectioning was done with sucrose-based dissection buffer and recorded in the standard ACSF as described in the **Electrophysiology and pharmacology** section above. Whole cell current-clamp recording of GCs was performed using an internal solution containing (in mM): 140 K-gluconate, 10 EGTA, 10 HEPES, 20 Na-phosphocreatine, 2 Mg-ATP, 0.3 Na-GTP, and 0.1 spermine adjusted to pH 7.3 and 310 mOsm with KOH. The membrane potential of the GCs was monitored in response to stimulation of the perforant path fiber with theta glass pipet connected to a stimulator (A-M System Model 4100). The stimulation protocol consisted of 10 sets of five distinct, 2 seconds, 10 Hz Poisson trains, delivered every 5 seconds. These input spike trains were generated using Matlab (R2019a, Mathworks) and followed a Poisson distribution with a similarity between trains set at R_input_ = 0.76 (average Pearson’s correlation coefficient with a bin window of 10 ms). The intensity of the stimulation was determined to obtain a probability of inducing a spike in response to the stimulus in the range of 30%-80%. The level of convergence or separation operated by the circuit was then calculated based on the comparison of the similitude of the inputs spike trains with the similarity of the recorded outputs spike trains. Multiple metrics were used to measure the similarity of the inputs spikes trains and the 50 recorded output traces (10 times 5 inputs). Each metrics used (i.e. R, NDP and SF) informs about distinct aspect of the spiking pattern such as firing rate, temporal coincidence of the spikes, “othogonalization” of the signal, and scaling factors; all representing putative neural codes (Madar et al., 2019b). Results were reported as pairwise comparison of the similarity of the inputs versus those of the outputs (e.g. NDPinputs versus NDPoutputs). Analysis of covariance (ANCOVA) were used to test statistical significance of the correlation comparison. Because all inputs were generated with a R_input_ = 0.76, the mean of the outputs for each metrics was also compared. In such cases, the statistical analysis was performed using a two-sample t-test. To analyze the firing pattern of the outputs, we calculated the firing rate as defined by the number of spikes per sweep and burstiness, the number of spike following a single stimulus. We further analyzed the transformation of the inputs spike trains by the circuit using additional metrics, compactness and occupancy. These metrics developed by Madar et al. are based on the binning of the spike trains and take into consideration the spikes distribution along the trains (“burstiness” of the trains); compactness takes into account the number of occupied bins in a trains whereas occupancy is defined by the number of spikes in the occupied bins. Dispersion metrics can then be calculated by performing pairwise comparison of the compactness and occupancy for the inputs and outputs spike trains of each recordings. Generated input trains and data analysis were performed using Matlab (R2019a, Mathworks), and original scripts were generously provided by Dr. Jones’s lab (https://github.com/antoinemadar/PatSepSpikeTrains).

### Statistics

All statistical analyses were performed using SPSS and Matlab (Mathworks). Except for pairwise analysis presented in figure 5, all data are presented as mean ± SEM. Data were analyzed with parametric test, including 2-tailed two-sample t-test or one-way ANOVA followed by Tukey post hoc analysis for comparisons of multiple samples. Probability distributions were compared using the Kolmogorov-Smirnov test. Sample sizes were not pre-determined but are similar to those reported by previous publications in the field (Hartzell et al., 2018; Soh et al., 2018). The digital numbers present within the histogram bars of all figures represent the number of animals per condition (top) and the number of biological replicates (bottom). Statistical details of experiments can be found in the figure legends. p values < 0.05 were considered statistically significant.

## Authors Contribution

Ting Feng, Conceptualization, Data curation, Software, Formal analysis, Investigation, Methodology, Supervision, Validation, Visualization, Writing—original draft, Writing—review and editing, Performed all electrophysiology, Analyzed all electrophysiology data, Performed immunohistochemistry, images acquisition, image analysis, virus production and injection; Christian Alicea, Performed image analysis; Vincent Pham, Performed spike trains analysis, Modified Matlab scripts; Amanda Kirk, Performed image analysis, Writing—editing; Simon Pieraut: Conceptualization, Data curation, Software, Formal analysis, Investigation, Methodology, Supervision, Validation, Visualization, Writing—original draft, Writing—editing, Funding acquisition, Project administration, AAV design and cloning.

## Acknowledgements

We thank Dr. Mathew V. Jones and Dr. Antoine D. Madar for their generous support, guidance and feedback on the recording and analysis for the spike train pattern separation experiment. We thank Dr. Jung Hwan Kim for his valuable comments and feedbacks on the manuscript. We thank Cynthia Lee, Aizel Nadonga, Catherine Bonuel and Blake Sieck for assistance with the mouse colony, genotyping and cloning of AAV vectors. Research reported in this publication was supported by Startup funds from the University of Nevada, Reno and by a grant from the NIGMS of the National Institute of Health under grant number P20 GM103650 to DM. Research reported here used the Cellular and Molecular Imaging Core facility supported by the National Institute of General Medical Sciences of the National Institutes of Health under grant number P20 GM103650.

---

**Figure S1.**
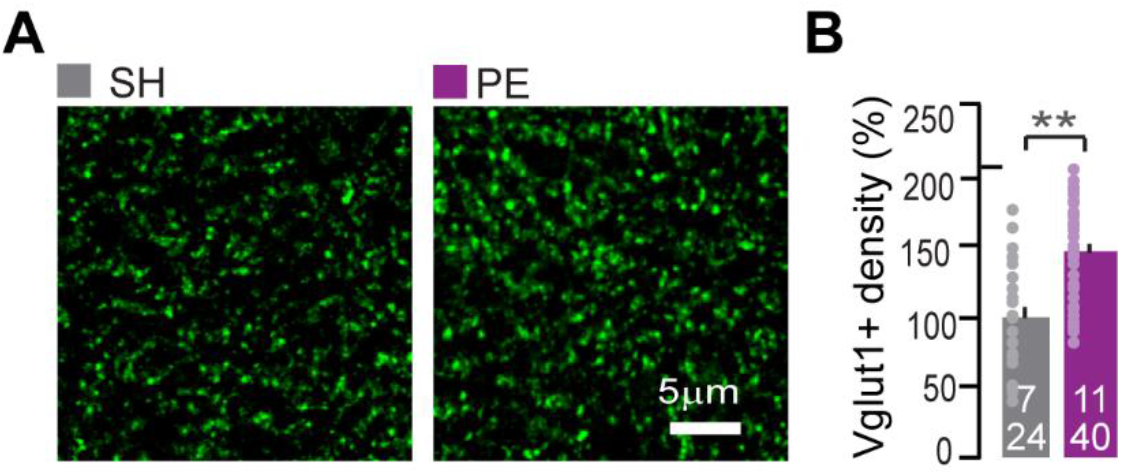
Experience-dependent increase of excitatory synaptic terminal. (**A**) Representative images of excitatory synaptic terminals labeled with antibodies against VGLUT1 (green) in the ML. (**B**) Quantification of VGLUT1 puncta in the ML of brain section from mice raised in SH (3-4 hemispheres from a total of 7 mice) and PE mice (3-4 hemispheres from a total of 11 mice). Graph bars represent mean ± SEM. PE data were normalized to SH control. Statistical comparisons were performed with a two-sample t-test (**: p<0.01).

**Figure S2.**
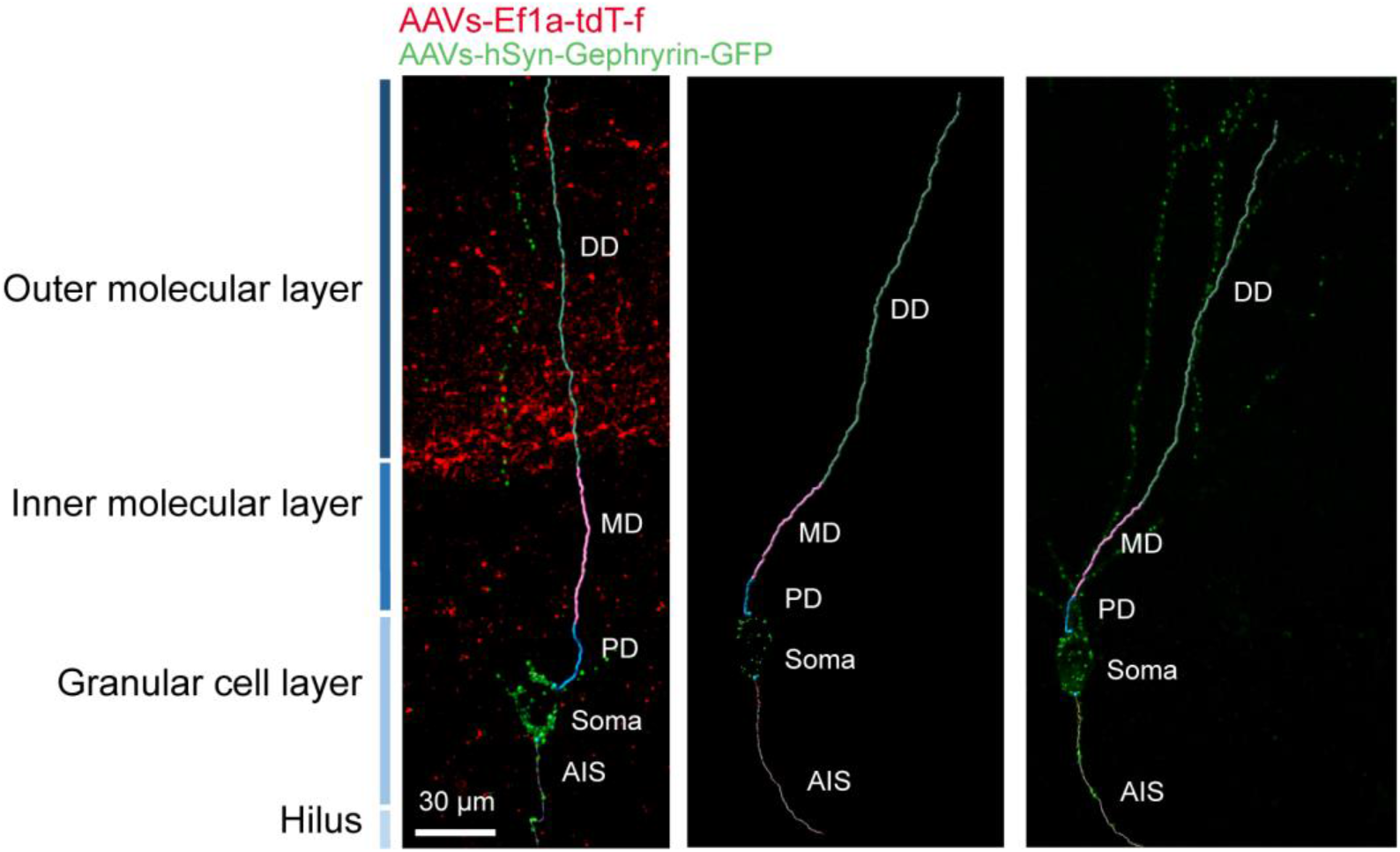
GC dendrites tracing and Gephyrin-GFP cluster analysis. Tracing and segmentation of GC dendrites with Imaris is performed using anatomical cue (GC nuclei) as well as with the help of the labeling of EC2 fibers with tdTomato to localize the separation between inner and the outer molecular layer (iML and oML). For this purpose, dual AAV injection was performed with AAV-hSyn-Gephyrin-GFP (green) and AAV-Ef1α-tdT-F. This method allows to distinguish the distal dendrite (DD), the medial dendrite (MD), the proximal dendrite (PD), the soma and the axons initial segment (AIS). The density of Gephyrin-GFP cluster per micrometers of dendrite were calculated within each segment (**Figure 2D**).

**Figure S3.**
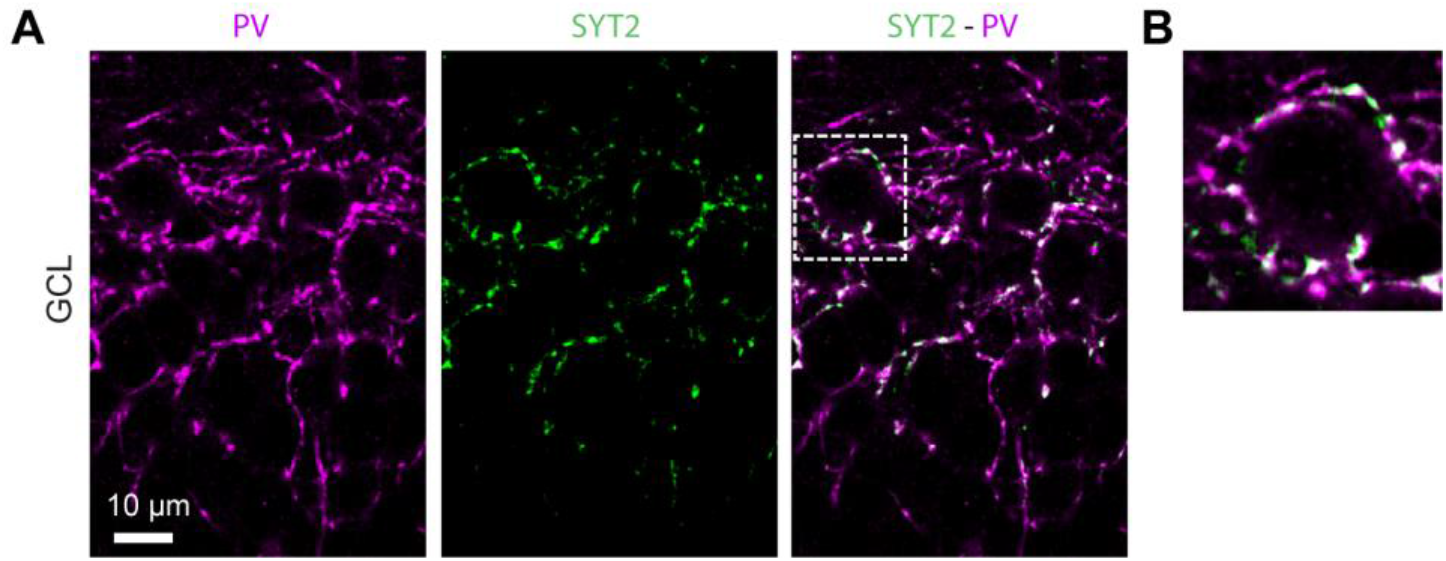
Syt2 is express in PV+ boutons in the GCL. (**A**) Representative images of the GCL stained with an antibody against PV (magenta) and an antibody against Syt2 (green). (**B**) Inset from the previous image showing that Syt2 accumulates in PV+ boutons labelled with the PV antibody and surrounding the mature GCs in the DG.

**Figure S4.**
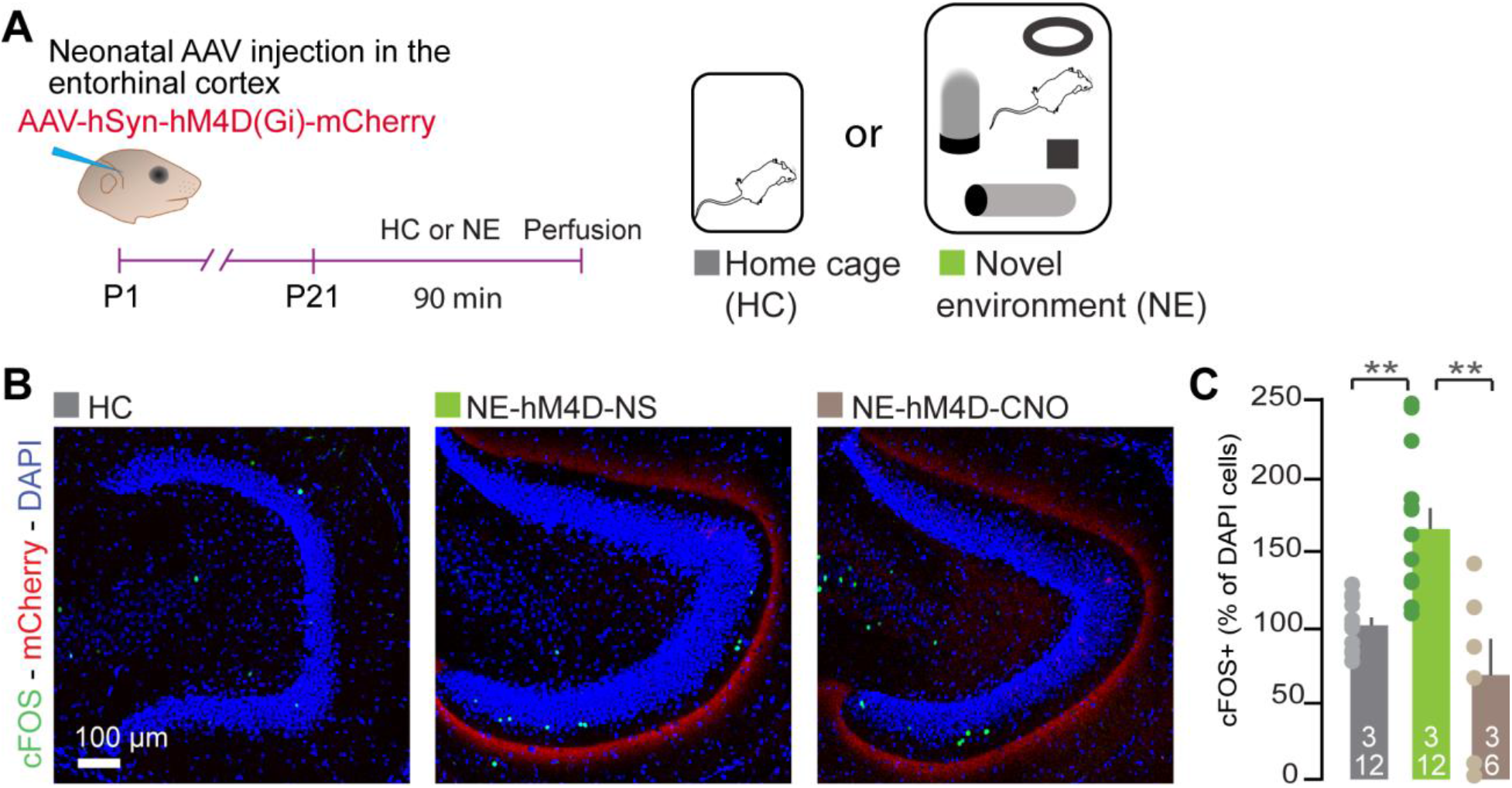
Inhibition of EC2 afferents using DREADD receptor hM4D(Gi). (**A**) Cartoon representing the experimental paradigm used to induce cFos expression in response to the exploration of a novel environment (NE) in mice injected with AAV-hSyn-hM4D(Gi)-mCHerry in the EC. (**B**) Representative images of the DG from brain sections labelled with an antibody against cFOS. The projections from EC2 neurons are labelled with mCherry and nuclei were labeled with DAPI. (**C**) Quantification of cFOS expressing cells in the GCL. Number is compared in sections obtained from HC group or mice expressing hM4D and placed in NE. Mice expressing hM4D received an IP with either normal saline solution (NE-hM4D-NS) or CNO (NE-hM4D-CNO). All data were normalized to the control HC values. Statistical comparisons were performed with an ANCOVA followed by Tukey HSD post-hock test (*: p<0.05). For each bars, the number of animals and the number of images used for the quantification are reported (a/n).

**Figure S5.**
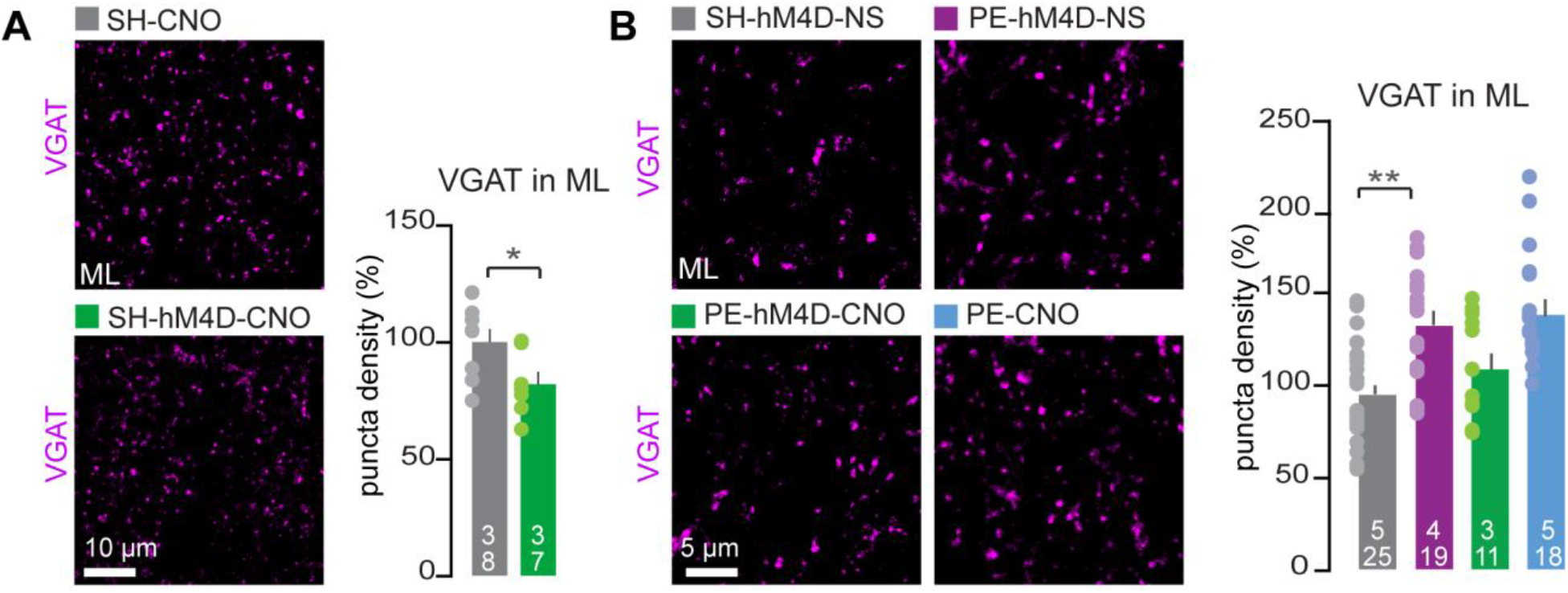
Density of presynaptic GABAergic terminals in the ML is regulated by the activity of the EC2 afferents. (**A**) Left, representative image of brain section stained with antibody against VGAT. Right, quantification of the VGAT puncta density in the ML of SH control mice (SH-CNO) or mice injected with AAV-hSyn-hM4D(Gi)-mCHerry in the EC (SH-hM4D-CNO), both received an IP with CNO. Graph bars represent mean ± SEM. All data were normalized to SH-CNO controls. Statistical comparisons were performed with a two-sample t-test (*: p<0.05). (**B**) Same as for (A) but chronic inhibition of the EC2 afferents was done in PE mice (PE-hM4D-CNO) and compared to values obtained from SH and PE mice also injected with the virus but who received NS treatment (SH-hM4D-NS and PE-hM4D-NS). An additional PE control group with no virus received the CNO injection (PE-CNO). Statistical comparisons were performed with an ANOVA (**: p<0.01) followed by Tukey HSD post-hock test. For each bars, the number of animals and the number of images used for the quantification are reported (a/n). All scale bar are in μm.

